# Human CD8^+^ T-cells Require Glycolysis to Elicit Effector Function

**DOI:** 10.1101/2020.02.05.935627

**Authors:** Scott E. Stimpson, Jing Chen, Brittney N. Newby, Ram Khattri, Harold D. Chapman, Thomas E. Angelini, Matthew E. Merritt, David V. Serreze, Clayton E. Mathews

## Abstract

Targeting human T-cell metabolism for modulating immune function requires an understanding of macronutrient utilization. Using metabolic inhibition during activation of human naïve CD8^+^ T-cells, we demonstrate blocking glycolysis or mitochondrial respiration prevents T-cell proliferation. However, after activation and differentiation, the metabolic program changes. Inhibition of glycolysis abolished cytotoxic T-lymphocyte (CTL) activity, whereas mitochondrial inhibition had no effect on CTL lytic function. Studies with uniformly labeled ^13^C-glucose confirmed CTL convert the majority of glucose to lactate. The role of glycolysis in CTL function was assessed using NOD models of Type 1 diabetes (T1D). Treatment of NOD models with a glycolysis inhibitor resulted in reduced and delayed T1D incidence and significantly preserved β-cell mass. We conclude glycolysis and mitochondrial ATP production are essential for efficient T-cell activation, but only glycolysis is essential for CTL lytic function. These data suggest targeting glycolysis in CTLs is a promising pathway to prevent T-cell-mediated autoimmunity.

---

Immune responses are critical for eliminating pathogens, but are tightly controlled to prevent unnecessary inflammation and autoimmunity. Recent studies provided evidence that activation of immune cells requires cellular metabolism. Adenosine triphosphate (ATP) is an important energy source produced through glycolysis and oxidative phosphorylation (OXPHOS)^1^. Though little is currently known about the metabolic requirements of human cytotoxic CD8^+^ T-cells or the macronutrients needed to fuel activation, differentiation, and effector functions, the potential disease impacts of targeting immune cell metabolism emphasizes the need for a better understanding of the energy requirements of T-cells during these processes.

Previous studies focused on metabolic control and requirements of mouse lymphocytes, specifically CD4^+^ T-cells. Sena *et al*^1^, demonstrated a major role for mitochondrial complex III reactive oxygen species (ROS) for both *in vitro* and *in vivo* IL2 production in CD4^+^ T-cell activation. The major glucose transporter, Glut1, has been shown to play a role in metabolic reprogramming to drive glycolysis in CD4^+^ T-effector (T_eff_) cells. In T_eff_, Glut1 was required for growth and expansion, but for regulatory T-cells (T_reg_), Glut1 knockdown had little effect^2^. Target of rapamycin (mTOR) also plays a distinct role in activation-induced metabolic and transcriptional control, which is important for T_reg_ activation and function. mTOR inhibition reduces T_reg_ suppressive capacity^3^. It has also been shown T_reg_ cells require mitochondrial complex III to maintain their regulatory gene expression and suppressive function^4^. Furthermore, the tricarboxylic acid cycle (TCA) is required for terminal effector function of T_H_1 cells through succinate dehydrogenase (complex II), but the latter activity suppresses proliferation of activated mouse T_H_1 CD4^+^ T-cells^5^.

Glycolysis in mouse CD8^+^ T-cells is required for effector function upon activation, but less important for proliferation and survival^6^. Partial rapamycin induced inhibition of mTORC1 in mouse CD8^+^ T-cells promoted memory formation; however, *Raptor* deletion caused a loss of mTORC1 function impairing CD8^+^ T-cell activation and effector differentiation^7^. A deficiency in Rheb (Ras homolog enriched in brain), a GTP-binding protein that activates mTORC1 within mouse CD8^+^ T-cells, resulted in an ineffective effector response^8^. In addition to ATP production, generation of mitochondrial ROS plays a crucial role in mouse T-cell activation, driving antigen-specific expansion^9^. These previous studies form a foundation for the current investigation.

The demonstration that T-cells require specific metabolic pathways suggests targeting these pathways can slow or halt CTL-mediated killing. Mitochondrial ATP production is essential for efficient activation and proliferation of human CD8^+^ T-cells. Here, we demonstrate activated cytotoxic lymphocytes (CTLs) do not require OXPHOS for effector function. In activated CTL, glycolysis is essential for target T-cell lysis. Further, treatment of NOD models of CD8^+^ T-cell mediated autoimmune type I diabetes (T1D) with the glycolysis inhibitor 2-deoxy-glucose reduced or prevented disease. This investigation is innovative as it identifies the important yet understudied metabolic mechanism controlling human CD8^+^ T-cell function and autoreactivity by examining cellular nutrient metabolism by CTL.

## Results

### Expansion of naïve human CD8^+^ T-cells requires glycolysis and OXPHOS

To better understand the macronutrient requirements of naïve human CD8^+^ T-cells for activation and expansion, we isolated such cells from young, healthy donors. Isolated naïve CD8^+^ T-cells were cultured in cRPMI and activated in the presence of αCD3/CD28 beads, with varying substrate and mitochondrial inhibitors (Figure 1a & 1b). Naïve CD8^+^ T-cells, under normal physiological concentrations of 5.5mM glucose, were able to expand 12-fold in the 9-day period^10^. However, using less efficient energy sources, such as 5.5mM galactose, expansion capacity dropped to 9-fold compared to 5.5mM glucose (p>0.05), indicating that reduced glycolytic ATP production is sufficient to sustain expansion (Figure 1c). There was a slight increase in expansion capacity with 11mM glucose and 5.5mM glucose supplemented with 10mM methyl succinate, but this was not statistically significant (p>0.05). However, expansion capacity was almost ablated in the presence of 2DG and was not able to be rescued by addition of pyruvate or the mitochondria complex II substrate, methylsuccinate (p<0.0003). When mitochondria of naïve CD8^+^ T-cells were inhibited using complex I inhibitor rotenone or complex III inhibitor antimycin A, expansion capacity halted and cells subsequently died off (p<0.0001). The conditions of Complex I or Complex III inhibition were not subject to rescue with methylsuccinate (Figure 1c).

**Figure 1.**
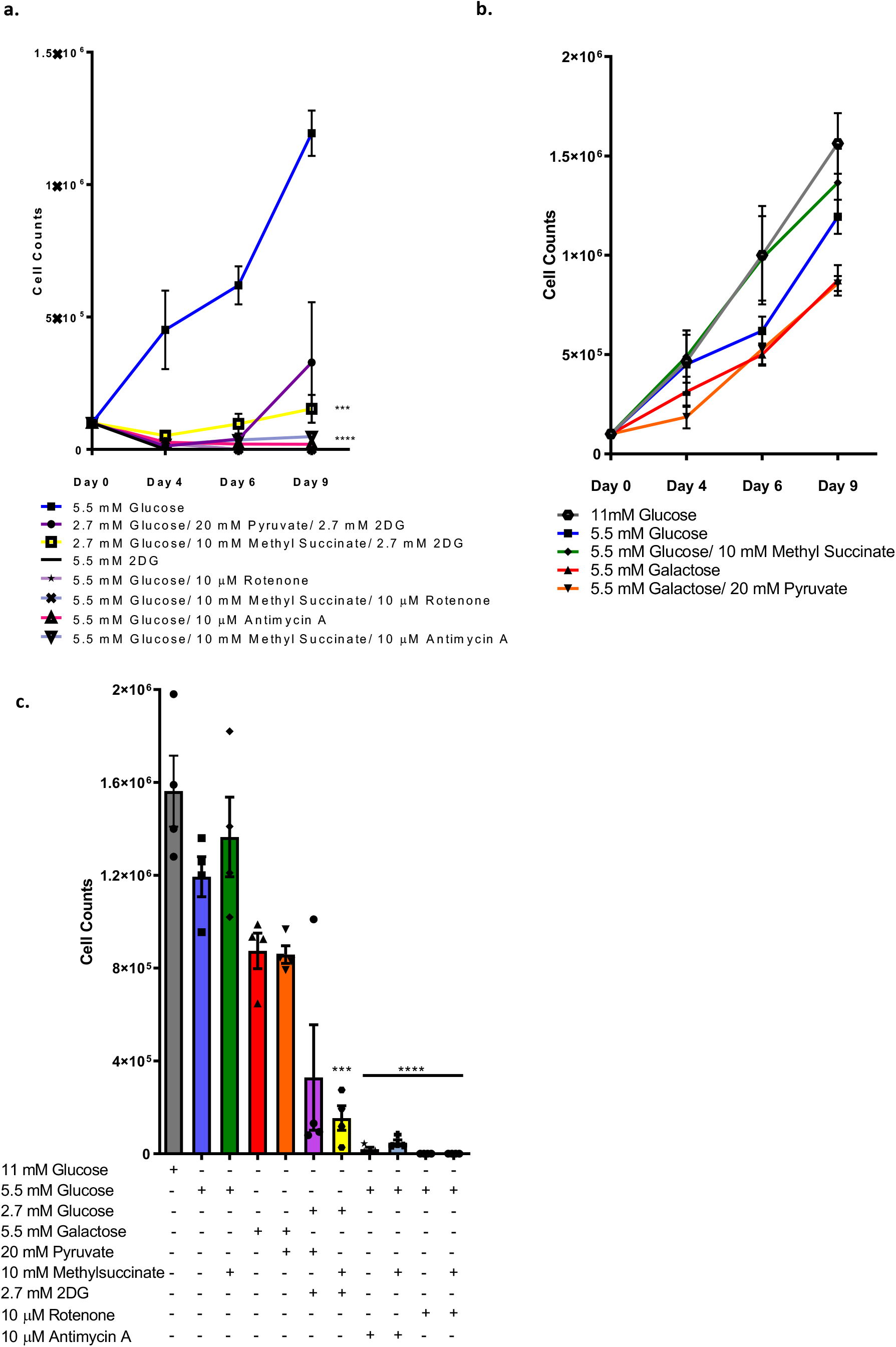
Human naïve CD8^+^ T cells require both mitochondrial and glycolytic ATP for activation and expansion. **a-b)** Line graphs displaying decreased cell numbers after 9 days of activation and expansion in all metabolic inhibitors with no rescue observed with the addition of metabolic substrates, and a reduced number of cells when galactose is the sole source of energy input. **c)** Day 9 cell count analyses indicate that that when naïve CD8^+^ T cells are in substrate limitation or excess, cells are unable to effectively activate and expand in comparison to 5.5mM glucose control. Mean ± SEM graphed. These data are compiled from 4 healthy donors. *** indicates *p*<0.0003 and **** *p* <0.0001 when compared to control.

Upon activation, expanded T-cells were assessed at Days 4 (**Figure S1**) and 9 (Figure 2) to determine the activation phenotype under each experimental condition. After polyclonal activation cells were assessed for memory or naïve phenotype marker expression (CD44 and CD62L). T-cells under normal glucose conditions were skewed to a more naïve phenotype, compared to T-cells that exhibited glycolytic inhibition and a memory phenotype (CD44 low, CD62L high). Furthermore, the expression of effector markers CD137, Granzyme B, and CXCR3 were also reduced in substrate limiting or inhibitory conditions (Figure 2c-2e). The results here show that human naïve CD8^+^ T-cells require both mitochondria and glycolytic energy production for successful activation and resulting clonal expansion. Previous studies demonstrated that T-cell expansion can be improved by certain metabolites being present upon antigen stimulation, and that memory T-cells have a more catabolic metabolic profile compared to that of effector cells which exhibit more anabolic metabolic profiles^5,11^. When we ablated mitochondrial activity *in vitro*, naïve CD8^+^ T-cells were unable to become activated or expand, resulting in T-cell death. However, naïve cells become activated and had a reduced expansion capacity when glycolysis alone was inhibited. Noteworthy here is that when glycolysis is inhibited by 2DG, the resulting expanded T-cells exhibited a memory phenotype, with decreased Granzyme B and CD137 expression. A reduction in the effector phenotypes of activated and expanded CD8^+^ T-cells would provide an avenue to decrease long-term autoimmunity such as pancreatic beta cell destruction underlying T1D development.

**Figure 2.**
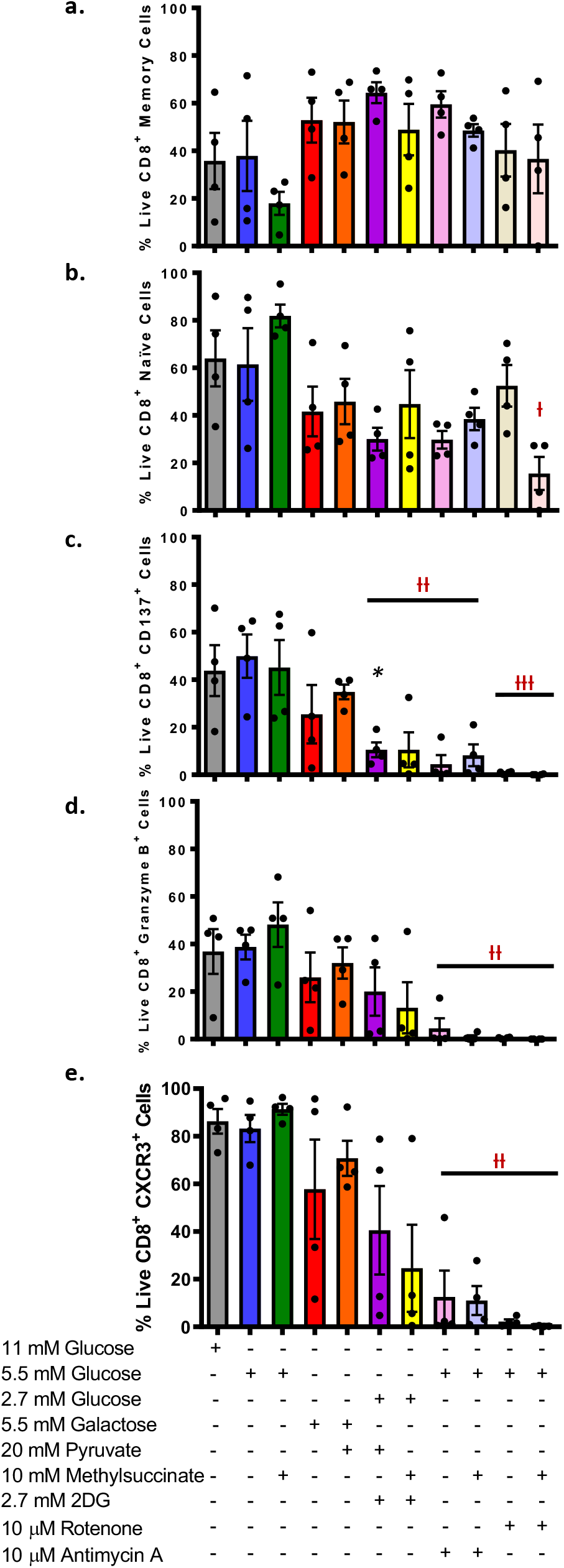
Human naïve CD8^+^ T cells exhibit decreased activation markers and Granzyme B levels in conditions of substrate limitation/excess after 9 days of expansion. **a-b)** Analyses reveals that the metabolic inhibitions forces cells into a more memory-like phenotype compared to that of glucose alone. **c)** Decreased expression of activation marker, CD137, occurs in all conditions of substrate imitation (no glucose) or glycolytic and mitochondrial inhibition. **d)** Decreased expression of cytolytic enzymes, Granzyme B, occurs in all conditions of substrate imitation (no glucose) or glycolytic and mitochondrial inhibition. **e)** Expression of CXCR3 is decreased when glycolytic and mitochondrial inhibition occurs. Mean ± SEM graphed. These data are compiled from 4 healthy donors. ᵻ denotes *p<0.01*, ᵻᵻ denotes *p<0.001 and* ᵻᵻᵻ denotes *p*<0.0001.

### Energy produced via glycolysis is essential for CTL effector function and release of Granzyme B

To determine the bioenergetic needs of CTLs for effector function and destruction of target cells, we co-cultured CTL-avatars with βlox5 cells, a human beta cell line kindly provided by Dr. Fred Levine (Sanford Children’s Health Research Center, Sanford-Burnham Medical Research Institute, La Jolla, CA). These co-cultures were used to measure antigen-specific CTL lytic activity in the presence of different macronutrient concentrations, as well as inhibitors of glycolysis and OXPHOS. Viability of IGRP-GFP CTLs (Figure 3a-3d) and βlox5 cells (Figure 3e-3h) under the described conditions were assessed to determine if the substrate or inhibitor conditions were toxic. With the exception of the complex I inhibitor rotenone, there were no statistically significant differences in toxicity to the βlox5 cells or CTLs when compared to the 5.5mM glucose control.

**Figure 3.**
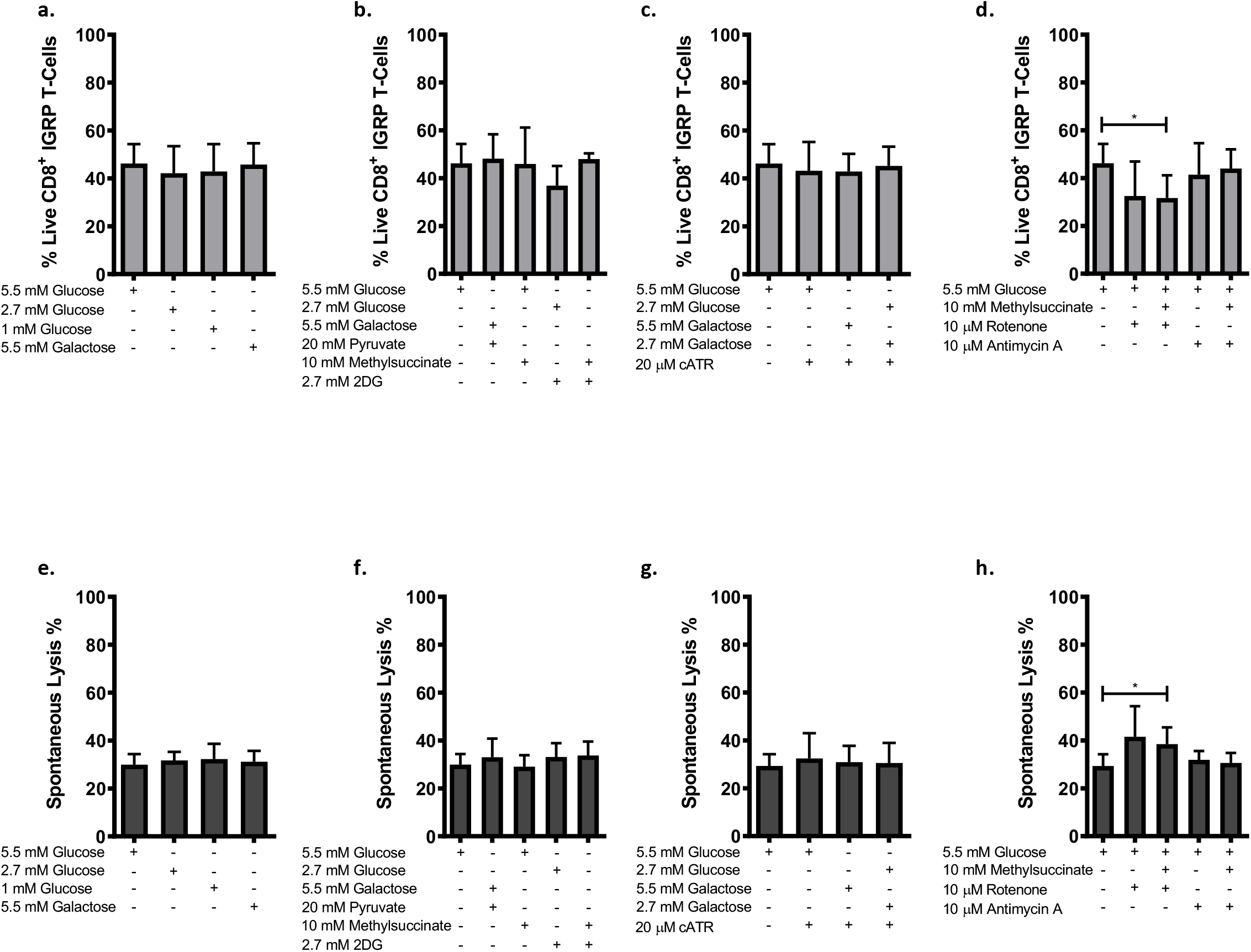
Metabolic inhibitors and substrate imitation/excess is not toxic to IGRP-CD8^+^ T cells after 16 hour culture. **a-d)** IGRP-specific CD8^+^ T cells were cultured for 16 hours in varying concentrations of glucose, galactose and metabolic substrates and inhibitors for 16 hours and cell viability determined by FACS analysis. **e-h)** βlox5 cells were cultured for 16 hours in varying concentrations of metabolic substrates ± inhibitors for 16 hours and cell viability determined by spontaneous chromium^51^ release (CML assay). Mean ± SEM graphed. * indicates *p* <0.05.

To determine the glucose-derived bioenergetic needs of CTLs for effector function and destruction of target βlox5 cells, we co-cultured IGRP-GFP CTLs with βlox5 cells at different effector to target ratios (0:1, 1:1, 5:1, and 10:1) for 16 hours (Figure 4). First, we observed the impact of glucose concentrations on CTL lytic activity. CTLs were able to carry out full effector functions in reducing glucose concentrations from 5.5 to 2.7mM, 1mM, or 0mM (Figure 4a). However, the addition of an equal molar concentration of glucose and 2DG resulted in complete ablation of CTL lytic function (p<0.0001, Figure 4b). CTL lytic function in the presence of 2DG was not rescued by the addition of the mitochondrial substrate’s pyruvate or methylsuccinate. This suggests that glycolysis is essential, but mitochondria are superfluous, for lytic activity.

**Figure 4.**
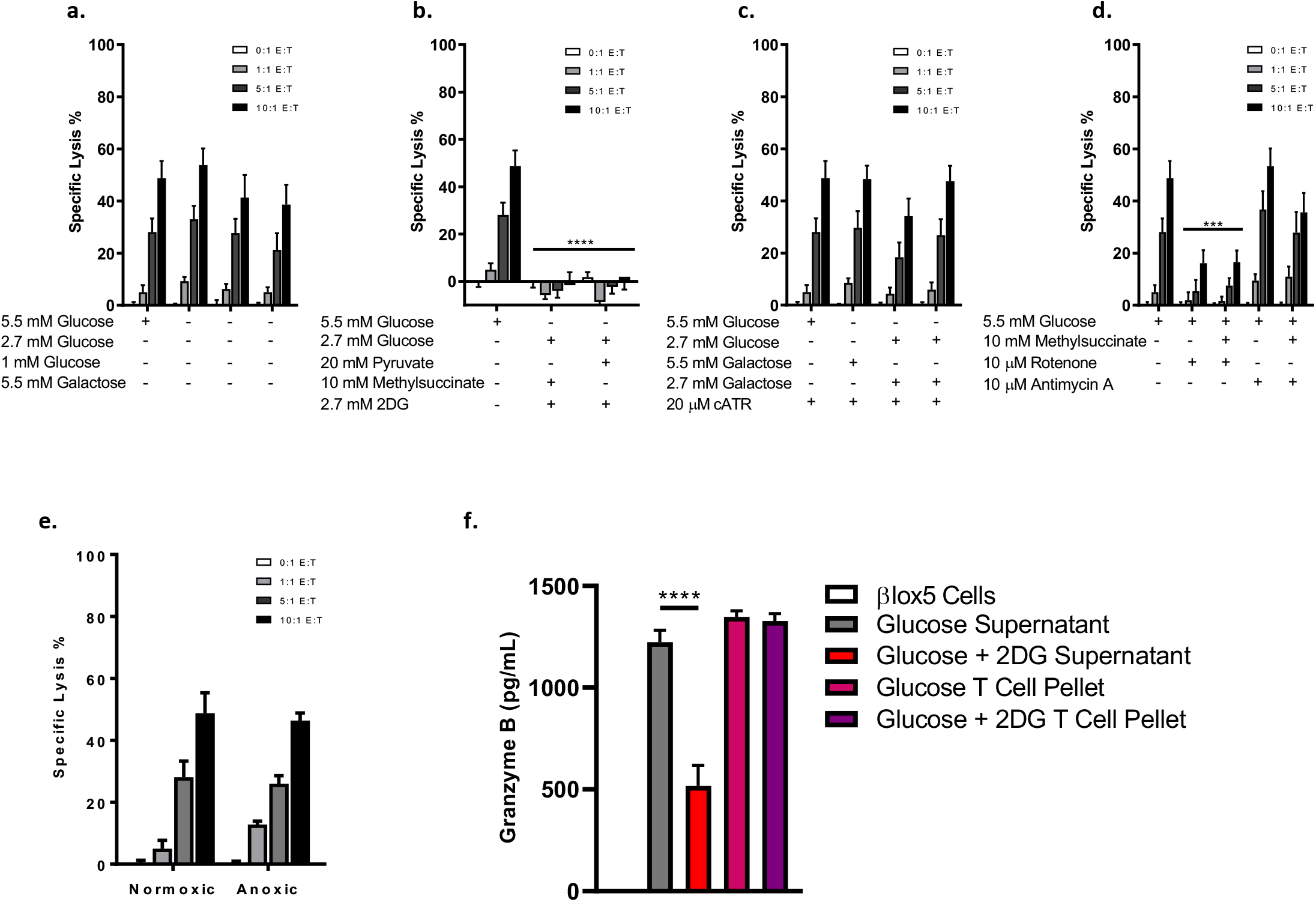
Two ATP from glycolysis is the bioenergetic requirement for CTL lytic function and not mitochondrial ATP production. βlox5 cells were primed with rhIFNγ (1000 IU/mL) for 24 hours. **a)** Cells were then co-cultured with IGRP-specific CD8^+^ T cells in varying concentrations of glucose or galactose for 16 hours. Specific lysis was determined using the cell mediated killing assay (CML) **b)** βlox5 cells were co-cultured with IGRP-specific CD8^+^ T cells in the presence of the glycolytic inhibitor 2DG, glycolytic substrate pyruvate and the mitochondrial electron transport chain complex II substrate, methylsuccinate, for 16 hours. **c)** βlox5 and IGRP-Specific CD8^+^ T cells were co-cultured in the presence of ATP translocase inhibitor, cATR, with either glucose or galactose. **d)** βlox5 and IGRP-Specific CD8^+^ T cells were co-cultured in the presence ETC complex I and III inhibitor, rotenone and antimycin A, and the complex II substrate methylsuccinate. **e)** βlox5 and IGRP-Specific CD8^+^ T cells were co-cultured in 100% nitrogen (anoxic) conditions for 16 hours. **f)** IGRP-specific CD8^+^ T cells were co-cultured with βlox5 16 hours in 5.5mM Glucose or 2.7mM Glucose and 2.7mM 2DG. Supernatants and T cell pellets were harvested and Granzyme B levels determined. Mean ± SEM graphed. *** denotes *p* <0.001 and **** *p*<0.0001. These data are compiled from 5 healthy donors.

In order to further elucidate the nature of mitochondrial input on lytic effector function of CTL, we utilized inhibitors of specific mitochondrial processes. Mitochondrial electron transport chain complexes I and II were inhibited utilizing rotenone and antimycin A, respectively. Blockade of complex I with rotenone resulted in reduced killing; however, this was due to the increased toxicity on the βlox5 cells, not the capacity of the CTLs to carry out their effector function (Figure 3h). The use of antimycin A on complex III did not alter the CTLs effector function. Likewise, addition of the complex II substrate, methylsuccinate, produced no alterations in activity (Figure 4d). ATP translocation out of the mitochondria was inhibited using cATR, with no decrease in killing observed (Figure 4c). This was also the same when glucose was replaced with galactose in conjunction with carboxyatractyloside (cATR). Furthermore, these data were confirmed when we assessed the ability of CTLs to destroy βlox5 cells in anoxic (100% nitrogen) conditions and discovered that, in little to no oxygen environment, CTLs can carry out their effector functions at the same level as normoxic conditions (Figure 4e). After confirming that glycolysis was essential for effector function, we determined if this was a result of reduced ability of the CD8^+^ T-cells to release granzyme B when co-cultured with βLox5 target cells. IGRP-GFP CD8^+^ T-cells that were co-cultured in the presence of 2DG had a statistically significant decrease (*p*<0.0001) in granzyme B levels present in the supernatant compared to that of the control (Figure 4f). Despite the reduction in granzyme B levels in the supernatant, both control and 2DG treated CD8^+^ T-cells had similar levels within the cell pellets, indicating that granzyme B was unable to be released during the co-culture period of those treated with 2DG, but the enzyme production is unaffected.

We have clearly shown that when mitochondria are inhibited at multiple ETC complexes and truncation of mitochondrial produced ATP occurs, antigen-specific IGRP-GFP CTLs effectively destroy beta cells with no reduction in viability of these effectors. In contrast, when we blocked glycolysis with 2DG, the CTLs were unable to elicit the destruction of the βlox5 cells, thus indicating the direct requirement of ATP produced via glycolysis. This result is strengthened by the inability of CTLs effector function to be rescued by the addition of pyruvate, the end product of glycolysis, directly, and by the ability of T-cells to carry out their effector function in anoxic conditions, which halt oxidative phosphorylation and highlight the sole requirement of glycolytic ATP. Additionally, CD8^+^ T-cells are unable to release granzyme B in the presence of 2DG, highlighting the needs of glycolytic ATP production for the exocytosis of granzyme B, which is essential in T-cell-mediated killing of target cells.

### IGRP-GFP CD8^+^ T-cells migrate in the presence of 2DG

Upon determining that 2DG can arrest CTL effector function, we tested if there was a decrease in T-cell motility resulting from inhibition of glycolysis. To assess the motility of IGRP-GFP CD8^+^ T-cells when cultured in 2DG, we used Matrigel to create a 3D culture system. IGRP-GFP CD8^+^ T-cells were placed into an 8 well glass chamber slide with primary human islets, with either 5.5mM glucose media or 2.7mM glucose and 2.7mM 2DG containing media, and the motility of T-cells was analyzed on a confocal microscope for 16 hours. Upon assessment of the acquired images, IGRP-GFP T-cells were able to move and interact with primary human islets in media that contained 2.7mM 2DG comparatively to 5.5mM glucose media (**Figure S4a and S4b**). Further imagine analyses (**Figure S4c, S4d, and S4e**) revealed that IGRP-GFP CD8^+^ T-cells in 3D culture are able to move at a similar velocity, had the same amount of displacement from their starting locations, and were able to interact with pancreatic β cells in the presence of 2DG. These data denote that T-cells are mobile when glycolysis is inhibited. In summary, CTL can migrate and interact with target cells, but lytic activity is inhibited (Figures 1, 2, and 4) when glycolysis is blocked.

### During co-culture with target cells CTL favor the conversion of glucose to lactate versus glutamate

Next, we determined the utilization of glucose in CTLs using ^13^C-NMR. IGRP-GFP CD8^+^ T-cells were co-cultured for 16 hours with βlox5 cells in the presence of uniformly labeled ^13^C glucose or ^13^C galactose with and without 2DG. Collected samples were analyzed for the formation and amount of ^13^C-lactate and ^13^C-glutamate produced using jHSQC NMR^12^. Analysis revealed that CTLs produce more lactate compared to glutamate (40% compared to 15%, Figure 5a) in the presence of βlox5 cells. There was a reduction in lactate and glutamate production when galactose was the sole energy substrate; however, CTLs still continued to produce more lactate than glutamate (Figure 5b). The addition of 2DG reduced both the amount of lactate and glutamate produced in both uniformly labeled glucose and galactose conditions, which was indicated by the absence of ^13^C-glutamate peaks in the jHSQC and ^1^D projection analyses (Figure 5c). Ultimately, activated and expanded CTLs break down glucose *via* glycolysis, shuttle this to lactate formation, and do not drive mitochondrial respiration as indicated by reduced glutamate labeling. Glutamate effectively reads out substrate selection for acetyl-CoA production^13^.

**Figure 5.**
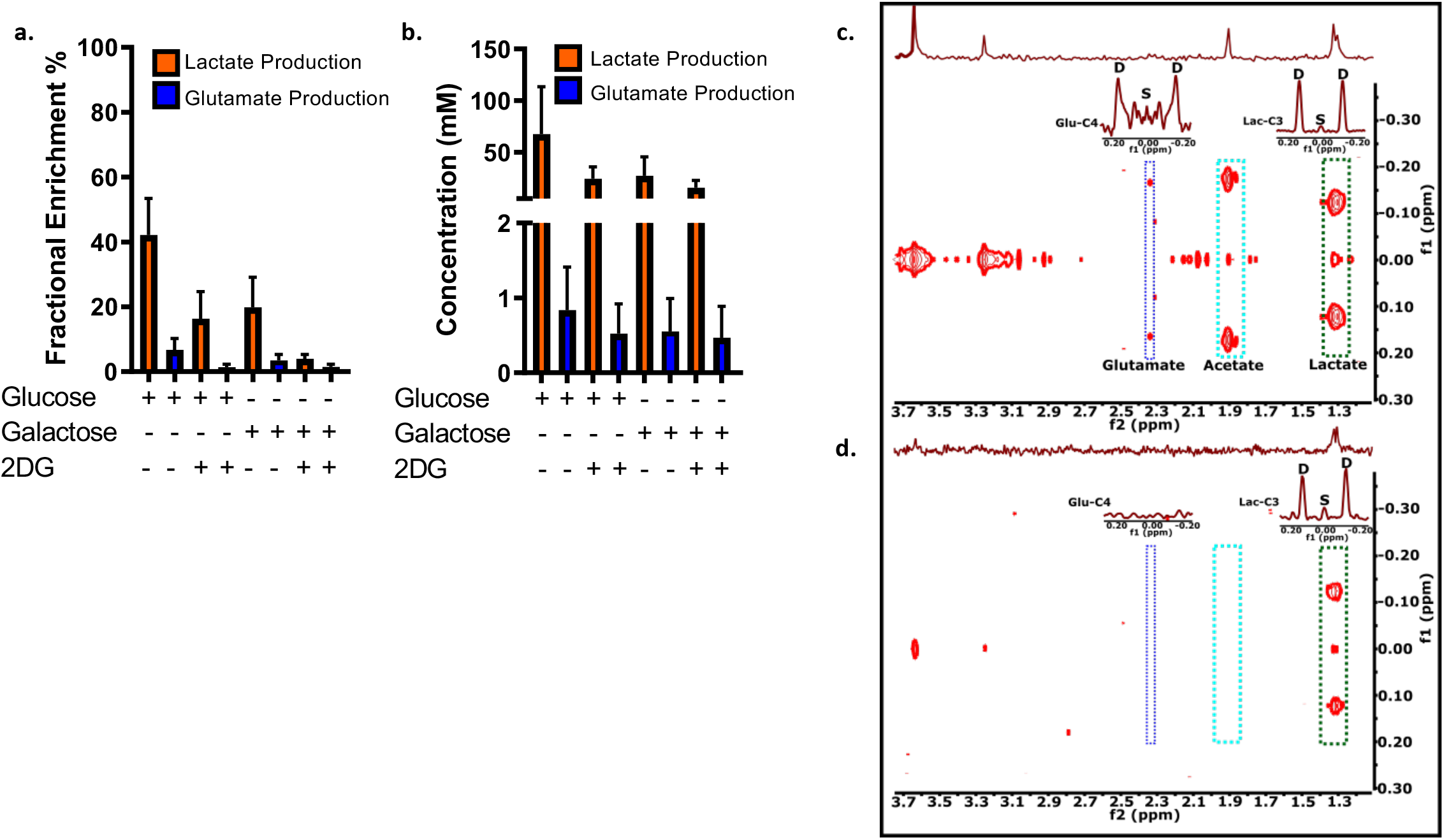
Reduced lactate and glutamate levels exhibited in the presence of U^13^C-Galactose and 2DG. βlox5 and CD8^+^ T cells were co-cultured at 30:1 in the presences of either U^13^C-Glucose, U^13^C-Galactose and 2DG **a)** U^13^C-labelled glutamate and lactate fractional enrichments were determined and analysed in T cell pellet and supernatants and combined. **b)** U^13^C-labelled glutamate and lactate concentrations were determined and analysed in T cell pellet and supernatants and combined. **c)** Representative jHSQC and ^1^D projections showing ^13^C labelling pattern for glutamate, acetate and lactate metabolites within T cell pellets that were labelled with U^13^C-Glucose **d)** Representative jHSQC and ^1^D projections showing ^13^C labelling pattern for glutamate, acetate and lactate metabolites within T cell pellets that were labelled with U^13^C-Glucose and 2DG. Glu-C4 represents C4 of glutamate and Lac-C3 denotes C3 of lactate. "S" and "D" denote singlet and doublet formations.

### 2DG delays diabetes onset in NOD/ShiLtJ and NOD.AI4 α/β-Thy1^α/b^ mice

In light of these data, we sought to understand CTL metabolism *in vivo* using a T1D model where metabolic activity of CD8^+^ T-cell destruction of pancreatic beta cells could be studied. 10-week-old female NOD/ShiLtJ mice were placed onto water bottles that contained either filter-sterilized water alone or water containing 35mM 2DG over a 20-week follow-up period. Diabetes incidence was monitored *via* tail vein blood glucose assessment, with a statistically significant increase in the survival of the 2DG treated mice compared to control water mice. An additional group of 10-week-old female NOD/ShiLtJ mice were treated with 35mM 2DG in their drinking water for 10 weeks and subsequently removed; diabetes incidence was monitored, with the majority of mice succumbing to diabetes after the removal of 2DG (Figure 6a). In addition, NOD.*Rag1*^*null*^.AI4αβTg (NOD.AI4) mice, a transgenic mouse model solely producing insulin-specific CD8^+^ T-cells were placed onto water bottles that contained either filter-sterilized water alone or water containing 35mM 2DG at 3 weeks of age. Spontaneous diabetes incidence was monitored every other day *via* tail vein blood glucose assessment. There was a marked delay in diabetes onset by 9 days in treated mice compared to control water mice (Figure 7a).

**Figure 6.**
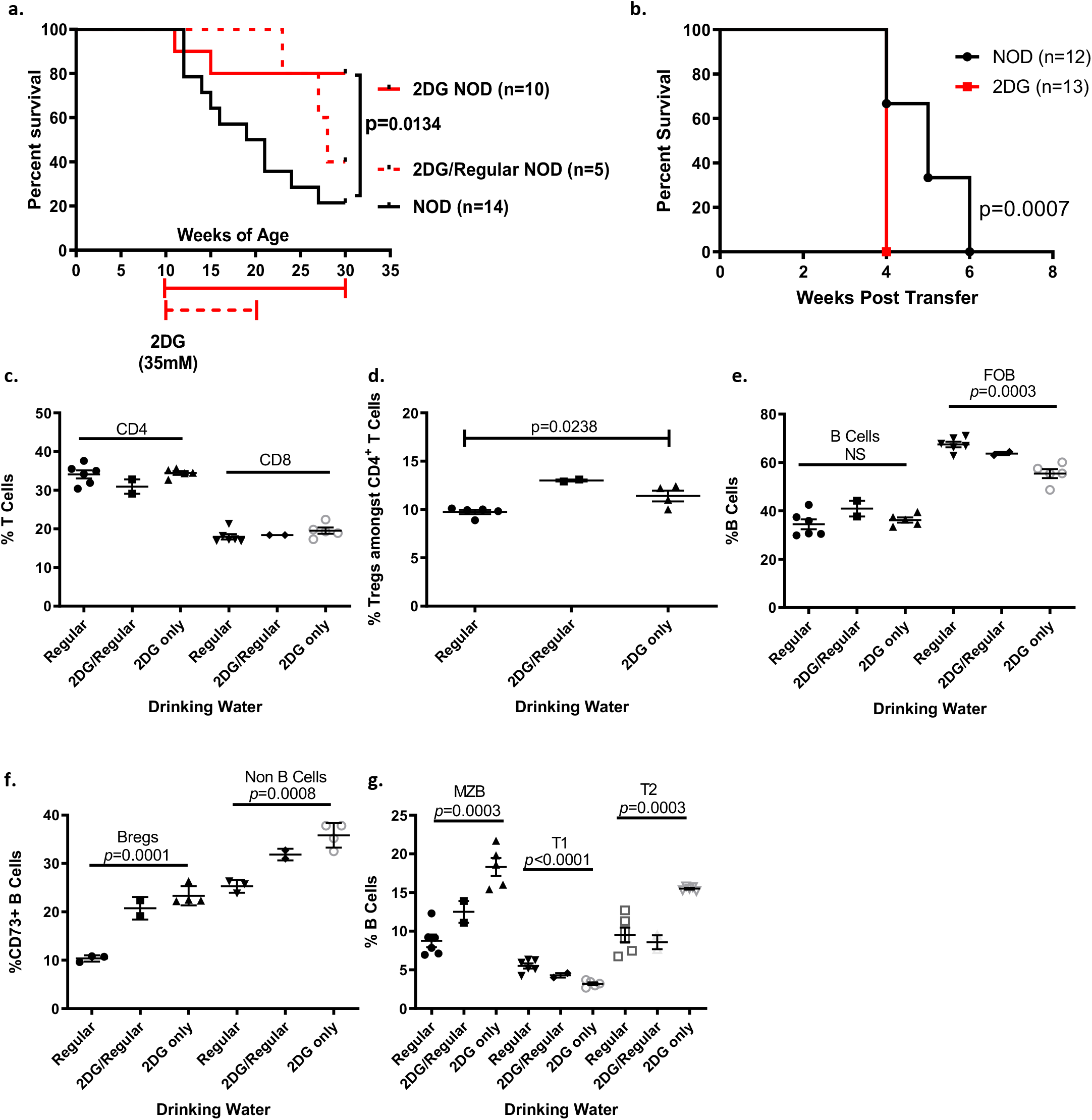
2DG administered in vivo delays the on-set of spontaneous diabetes in NOD/shiLtJ and NOD.AI4αβ mice. **a)** 10-week old female NOD/shiLtJ mice were treated with 35mM 2DG in their drinking water for 20 weeks and diabetes incidence monitored with a statistically significant increase in survival of the 2DG treated mice compared to control water mice. Red-dashed line indicates 10-week old female NOD/shiLtJ mice that were treated with 35mM 2DG in their drinking water for 10 weeks which was subsequently removed and diabetes incidence monitored. **b)** Splenocytes from 2DG treated or untreated NOD/ShiLtJ mice were adoptively transferred into NOD.CB17-Prkdcscid/J mice and incidence monitored post transfer. **c-e)** Total CD4^+^ and CD8^+^ T cells, CD4^+^ Tregs and B cells and follicular B cells were analysed in NOD/ShiLtJ mice that were placed onto regular drinking water, treated water with 35mM 2DG or treated for 10 weeks with 2DG and were placed back onto regular water. **f-g)** CD73^+^ Breg and non-B cells, and marginal zone B, T1 and T2 B cell subsets were analysed in NOD/ShiLtJ mice that were placed onto regular drinking water, treated water with 35mM 2DG or treated for 10 weeks with 2DG and were placed back onto regular water.

**Figure 7.**
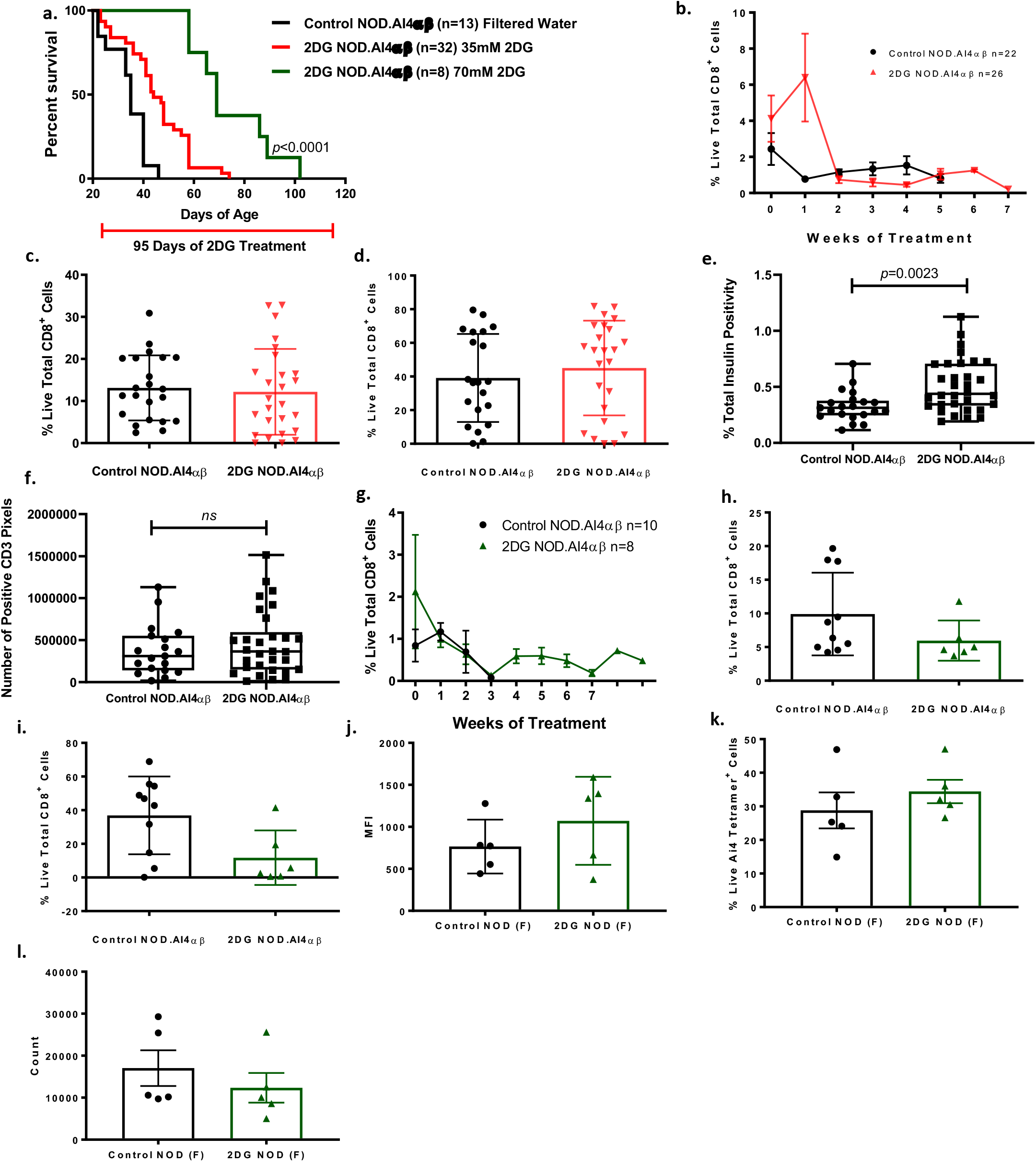
2DG administered in vivo delays the on-set of spontaneous diabetes but does not alter total CD8^+^ T cell numbers or halt proliferation in NOD.AI4αβ mice. **a)** 3-week old NOD.AI4αβ mice were treated with 35mM and 70mM 2DG in their drinking water and incidence of diabetes monitored until mice reached a blood glucose level of >600mg/dL on two consecutive days. Analysis revealed a statistically significant increase in the delay of diabetes onset in 70mM 2DG treated mice compared to that of untreated mice. **b)** Weekly cheek bleeds were carried out and total CD8^+^ T cells monitored in control and 35mM treated mice until diabetes on-set was reached. **c-d)** Upon reaching diabetes on-set mice were sacrificed and total CD8+ T cells analysed in the spleen and pancreatic lymph node respectively. **e)** Image analysis was carried out to determine total insulin positive staining in pancreatic sections in control and 2DG NOD.AI4αβ mice. Analysis revealed a statistically significant increase in insulin positive islets compared to control. **f)** Image analysis was carried out to determine total CD3 positive staining in pancreatic sections in control and 2DG NOD.AI4αβ mice. Analysis revealed no statistically significant difference in CD3 positive cells compared to that of control mice. **g)** Weekly cheek bleeds were carried out and total CD8^+^ T cells monitored in control and 70mM treated mice until diabetes on-set was reached. **h-i)** Upon reaching diabetes on-set mice were sacrificed and total CD8+ T cells analysed in the spleen and pancreatic lymph node respectively. **j)** Pancreatic lymph nodes from sub-lethally irradiated 70mM treated NOD/ShiLtJ were removed 3 days post injection with CTV labeled NOD.AI4αβ splenocytes and Ai4 tetramer^+^ cells analysed for Ki67 expression. **k)** Live Ai4 tetramer^+^ cells were analysed with no difference observed in treated mice pancreatic lymph nodes compared to that of control mice. **l)** No difference was observed in cellular numbers obtained from the pancreatic lymph nodes of 70mM treated mice compared to control mice. Mean ± SD graphed.

After determining that 35mM 2DG delayed the onset of diabetes, we wanted to determine whether 35mM 2DG treated NOD.AI4 mice had insulin preservation within the islets compared to that of the control. Analysis revealed a statistically significant retention (*p*=0.0023) of insulin positivity in 2DG treated mice (Figure 7b). Additionally, there was no difference in the amount of CD3 positive cells within the pancreatic sections of 35mM 2DG treated NOD.AI4 (Figure 7c). These data clearly show, and correlate with the above human data, that CD8^+^ T-cells are able to migrate and traffic into the pancreas, but are unable to illicit effector functionality, thus preserving insulin producing beta cells.

Upon diabetes onset, mice were euthanized; spleens were harvested from NOD/ShiLtJ mice, and peripheral blood, pancreatic lymph nodes, and spleens were harvested from NOD.AI4 mice. NOD/ShiLtJ mouse spleens were analyzed for total CD4^+^ and CD8^+^ T-cells, CD4^+^ Tregs, B and follicular B-cells (Figure 6c-6e), with no difference observed between the total CD4^+^ and CD8^+^ T-cell frequencies; however, Tregs and follicular B-cells were significantly altered in the 2DG treated mice compared to those treated with regular drinking water (*p*=0.0238 and *p*=0.003, respectively). Further analyses revealed significant changes in the frequencies of CD73^+^ B-cell and non-B-cell populations, both being significantly increased in the 2DG treated groups, as well as significant changes within marginal zone B-cells, T1, and T2 B-cell subsets (Figure 6f and 6g). Total CD8^+^ T-cells (Figure 7d-7f), as well as naïve, memory, or effector phenotypes, were also analyzed (**Figure S2**) from NOD.AI4 mice. There were no differences displayed in the naïve, memory, or effector populations of NOD.AI4 specific CD8^+^ T-cells treated with 35mM 2DG compared to the control mice.

With a greater effect seen in the NOD/ShiLtJ mice compared to the NOD.AI4 mice, we wanted to determine if increasing the concentration of 2DG in the drinking water of mice would alter the outcome in the NOD.AI4 model. We treated starting at 3-weeks of age NOD.AI4 mice with 70mM 2DG and monitored for them for spontaneous diabetes onset. The delay in diabetes onset was drastically significant when increasing the amount of 2DG, with NOD.AI4 mice surviving out to 37 ± 11 days (*p*<0.0001). However, there was no significant difference in the phenotype of AI4 CD8^+^ T-cell populations when compared to the control mice (Figures 7g-7i and **S3**). These *in vivo* data reflect the results observed *in vitro*, with CD8^+^ T-cells being unable to carry out their effector function effectively, resulting in delay of pancreatic beta cell killing as indicated by prolonged blood glucose stabilization.

### Diabetogenic NOD.AI4 CD8^+^ T-cells can proliferate in 2DG treated mice

With 70mM 2DG significantly delaying the onset of diabetes incidence, we investigated the ability of the NOD.AI4 CD8^+^ T-cells to proliferate in 70mM 2DG treated mice. Starting at 6-weeks of age NOD/ShiLtJ mice were treated with 70mM 2DG in their drinking water or received filter-sterilized water for 2 weeks prior to receiving a sublethal 750R irradiation dose. After irradiation, 3-week-old NOD.AI4 splenocytes were stained with a cell tracking dye, Cell Trace Violet (CTV), and injected into the tail vein of the control and treated NOD/ShiLtJ mice. After 3 days, peripheral blood and pancreatic lymph nodes (PLN) were harvested from the control and 2DG NOD/ShiLtJ mice, and AI4 CD8^+^ T-cells were assessed for the presence of CTV. The isolated AI4 tetramer^+^ cells contained no non-diluted proliferation dye, indicating that these cells can proliferate in the presence of 70mM 2DG, with further analyses revealing that there was no difference between the control and treated NOD.AI4 CD8^+^ T-cells expression of Ki67 or cells present in the PLN (Figure 7j-7l).

To further test that 2DG treated cells are able to confer disease, splenocytes were isolated from 2DG treated and untreated NOD/ShiLtJ mice and adoptively transferred into lymphocyte deficient NOD.CB17-*Prkdc*^*scid*^/J recipients, and diabetes incidence was monitored (Figure 6b). The adoptive transfer of splenocytes from 2DG treated NOD/ShiLtJ mice were able to confer disease in NOD.CB17-*Prkdc*^*scid*^/J, indicating that the effects of 2DG are only transient, coinciding with our *in vitro* data. With all mice succumbing to diabetes, these data demonstrate that long-term treatment of 2DG can slow down, but is unable to completely stop, beta cell destruction. In addition, AI4 CD8^+^ T-cells can still proliferate within 2DG treated mice post-adoptive transfer. Contrary to the above human data, the treatment of 2DG did not alter the phenotypes of the mouse AI4 specific CD8^+^ T-cells when compared to control groups (**Figures S2 and S3**), but did, however, alter the CD4^+^ Treg and B cells populations within NOD/ShiLtJ mice. This potentially highlights a distinct difference in responses of glycolytic inhibition of mouse T-cell subsets in comparison to humans. Further investigation is warranted, but this strengthens the need for more human studies to be carried out in T1D to ensure that we are effectively understanding patient relevant *in vivo* phenotypes.

## Discussion

This study opens new areas of investigation on how CD8^+^ T-cells can be targeted to prevent autoimmunity. Studies in disease models of lupus have previously shown that 2DG treatment of mice reduced the biomarkers and disease-associated changes through normalization of T-cell metabolism^14^. In T1D, studies using murine models have indicated that T-cell mitochondrial dysfunction may participate in the pathogenesis^15^. We have also more recently discovered that T-cells from patients with T1D, exhibit mitochondrial inner membrane hyperpolarization (MHP) when compared to controls^15^. Further, mitochondrial respiration and glycolysis are dysregulated in T-cells from T1D patients. T-cells from patients with T1D display aberrant metabolism and bioenergetics resulting in pathogenic functional activity including sustained activation, heightened effector function, and resistance to normal mechanisms of peripheral deletion and immune regulation. Our data here suggest that lymphocyte mitochondria and metabolic pathways are a promising and novel target for reversing autoimmunity in the prevention and cure of T1D.

The enclosed investigations provide novel information by specifically looking at human CD8^+^ T-cell metabolic requirements. These studies contribute and extend our current knowledge of T-cell metabolic requirements. Greater insight into how human CD8^+^ T-cells utilize energy substrates will provide essential tools for other autoimmune disorders, in addition to T1D. Recent studies have shown how type 1 interferons alter the signaling pathways in both a mitochondrial dependent and independent manner^10–16^, thus indicating an avenue of investigation to determine the effect of type 1 interferons in the inflammatory microenvironment upon human CD8^+^ T-cell metabolism^10^. With the ability to trace macronutrient usage during T-cell activity, we could alter metabolism to better understand how CD8^+^ T-cells require ATP and are able to better mimic the islet microenvironment, as well as providing greater insight for cancer or other immunotherapies and autoimmune diseases.

## Material and Methods

### Study subjects

Peripheral blood mononuclear cells (PBMC) were obtained from normal, healthy donors through the University of Florida Diabetes Institute (UFDI) study bank or from Leukopak samples obtained from LifeSouth Community Blood Centers (Gainesville, FL). Age demographic for female donors (n=6) were 15-26 years of age, and male donors (n=5) were 20-25 years of age. Ethical approval was obtained for these studies from the University of Florida Institutional Review Board (UF-IRB). Experiments with human materials were carried out under UF-IRB study numbers IRB201400703 and IRB201400933. Primary human islets were obtained from the NIH-funded Integrated Islet Distribution Program (IIDP). Donor islet Research Resource Identification (RRID) numbers used in this study were: SAMN08912443. Detailed islet donor information is outlined in **Supplemental Table 1**.

### Animals

All mice used in this study were housed in specific pathogen-free facilities. These studies were approved by Institutional Animal Care and Use Committees at our institutions and carried out under the protocol numbers 201509006 (University of Florida), 201809006 (University of Florida), or 99063 (The Jackson Laboratory).

### Creation of antigen specific CD8^+^ T-cell avatars

Antigen specific human CD8^+^ T-cells were produced as previously described^10,16^. All cultures were carried out in complete RPMI [cRPMI recipe: RPMI 1640 (Cellgro, Manassas, VA), supplemented with 10% FBS (HyClone, Fisher Scientific, Pittsburgh, PA), 1% MEM non-essential amino acids (Cellgro), 1% penicillin-streptomycin (Gemini Bio-Products, West Sacramento, CA) solution, 1% Glutamax (Cellgro), 1% sodium pyruvate (Cellgro), 0.004% 2-mercaptoethanol (Cellgro), and 10 mM HEPES (Cellgro)]. Briefly, naïve CD8^+^ T-cells were isolated from an enriched PBMC fraction and activated with αCD3 and αCD28 beads (Dynabeads, Invitrogen) for 48 hours. After 48 hours, these cultures containing bead activated CD8^+^ T-cells were transduced with a lentivirus containing a polycistronic construct encoding the HLA-A02*01 restricted Glucose-6-Phosphatase Catalytic Subunit 2 (IGRP) reactive T-cell receptor (TCR α/β chains from Clone 32), as well as green fluorescent protein (GFP) or the HLA-A02*01 restricted Melanin-A antigen recognized by T-cells (MART-1) reactive T-cell receptor (TCR α/β chains), as previously described^10,17,18^. Post-transduction, cells were cultured for an additional 7 days. On Day 9, cells were sorted for GFP positivity using a BD FACSAria III (BD Biosciences).

### Naïve CD8^+^ Expansion

Leukopak-derived peripheral blood mononuclear cells (PBMCs) purchased from LifeSouth Community Blood Centers were debulked with RosetteSep CD8^+^ enrichment kits (Stem Cell Technologies), and then separated into an enriched fraction of PBMC using a Ficoll gradient. Naïve T-cells (CD8^+^CD45RA^+^CD45RO^−^) were subsequently FACS purified using anti-CD8, as well as anti-CD45RA^+^ (Naïve) and CD45RO^−^ (memory) antibodies on a BD FACS Aria III. Purified naïve CD8^+^ T-cells were cultured at a density of 100,000 cells per well using cluster 48 plates in cRPMI media containing metabolic inhibitors and substrates combinations (5.5 & 2.7 mM glucose, 5.5 & 2.7 mM galactose, 2.7 mM 2-deoxy-D-glucose, 10mM dimethyl succinate, 20 mM pyruvate and 10µM antimycin A and rotenone). Cells were subjected to activation using αCD3 and αCD28 beads (Dynabeads, Invitrogen) at a 1:1 ratio. Cultures were supplemented with human IL-2 (100 U/mL, Prep Rotech USA) and additional 1mL of media every other day. Cells and supernatant were collected after 3, 6, and 9 days of expansion. Cells were analyzed by flow cytometry to assess activation (anti-CD44, CD69, and CD62L) and T-cell subset polarization (anti-CXCR3) using a BD LSR Fortessa (BD Biosciences, USA). Antibody sources and clones detailed in **Supplemental Table 2**.

### CML Assay

All cultures were carried out in complete DMEM [cDMEM recipe: low glucose (1mg/mL) DMEM (Cellgro, Manassas, VA), supplemented with 10% FBS (HyClone, Fisher Scientific, Pittsburgh, PA), 1% MEM non-essential amino acids (Cellgro), 1% penicillin-streptomycin (Gemini Bio-Products, West Sacramento, CA) solution, 0.02% BSA (Sigma, St. Louis, MO), and 15 mM HEPES (Cellgro)] ^19^. βlox5 cells were used as targeT-cells in the CML assays, and IGRP-GFP CTL avatars are effector cells, as previously described^10^. For hypoxic CML assays, βlox5 and CD8^+^ IGRP-GFP T-cells were plated as described above and cells were under hypoxic conditions in a 100% N_2_ atmosphere. The chamber was sealed and placed in a 37 °C incubator for the remaining 16 hours. Percent specific lysis was calculated as follows: [(Experimental release - Spontaneous Release) / (Experimental Release + Experimental Lysate) - Spontaneous Release] x 100^19–21^.

### Live/Dead Assay

CD8^+^ IGRP-GFP T-cells were plated at 200,000 per well in round bottom 96-well plates in the media containing the following substrates or metabolic inhibitors: 5.5 or 2.7 mM D-glucose, 5.5 & 2.7 mM D-galactose, 2.7 mM 2-deoxy-gucose, 10mM methyl-succinate, 20 mM pyruvate, 20 µM carboxyatractyloside (cATR), 10µM antimycin A, 10µM rotenone. T-cells were stained with Live/Dead Far Red for 30 minutes, subsequently washed, and then stained with anti-CD8 (PE) for an additional 30 minutes. T-cells were then washed and analyzed by flow cytometry (BD Accuri C6).

### *In vitro* movement of T-cells

Primary human islets were stained with 25nM Tetramethylrhodamine Methyl Ester Perchlorate (TMRM) for 30 minutes prior to culturing. Islets were handpicked and placed 10 per well into an 8-well chambered cover glass slide [Labtek glass chamber slide no.1.] coated for 30 minutes with 100µL of Matrigel (Corning, USA) at 37 °C, 5% CO_2_. CD8^+^ IGRP-GFP T-cells, 100,000 cells per well, were added in minimal volume, next to the plated islets. A further 100µL of Matrigel was slowly added on top and placed in 37 °C, 5% CO_2_ for 30 minutes to allow Matrigel to solidify. 300µL (containing 5nM TMRM) of 5.5mM 2DG or control DMEM media was layered on top. Z-stack images were acquired using the Zen 2.1 and collected using a Carl-Zeiss 710 LSM confocal scanning microscope in a humidified environment maintained at 37 °C, 5% CO_2_. Cell movement was measured and calculations performed, as previously described^22^, by analyzing maximum intensity time-lapse micrograph containing czi files in ImageJ.

### Granzyme B ELISA Assay

Supernatants and cell pellets were collected from the interaction between IGRP-GFP CD8^+^ T-cells and βlox5 cells at a 10:1 ratio and were evaluated using a granzyme B ELISA kit (BMS2027; ThermoFisher Scientific, USA.). The standard curve was produced to determine the granzyme B concentration. In brief, samples were collected after a 16-hour co-culture in either 5.5mM glucose or 2.7mM glucose and 2.7mM 2DG. The control groups had βlox5 cells alone. Cell pellets were lysed using PBS and rapid freeze-thaw cycles. Subsequently, the samples were incubated with an antibody-coated plate (from the ELISA kit) for further detection. All the procedures were performed according to the manufacturer’s protocols.

### NMR Sample Preparation

Samples were prepared by seeding βlox5 cells at a concentration of 1 × 10^6^ cells per well 24 hours prior to the addition of non-antigen specific MART-1 CD8^+^ T-cell avatars. T-cells were added at an effector to target ratio of 30:1 and co-cultured with the seeded βlox5 cells in either media containing 5.5mM U^13^C labelled glucose or 5.5mM galactose. In other studies, 2.7mM U^13^C-glucose or 2.7mM U^13^C-galactose was added in combination with 2.7mM 2DG and co-cultures continued for 16 hours. Cellular pellets and supernatants were harvested and snap frozen until sample analyses were carried out.

Metabolites present in the cells and supernatant were isolated via perchloric acid (PCA) extraction. Ice cold 6% (v/v) PCA was added to the cells and the mixture was homogenized using a FASTPREP-24 (MP Biomedicals, Solon, Ohio, USA) and centrifuged at 4 °C. The resulting supernatant was transferred into a new vial and neutralized with 5M potassium hydroxide. The neutralized solution was further centrifuged to remove insoluble potassium perchlorate. The supernatant was dried by lyophilization (Thermo-Scientific, Dallas, USA). The dried powder was dissolved in 200μL double distilled water, and the pH was adjusted using sodium hydroxide and hydrochloric acid. Excess salts were removed by centrifugation and the samples were lyophilized again. Supernatant samples were treated in the same manner, except no homogenization was required. The final NMR sample was prepared by re-suspending the freeze-dried sample in 90% (v/v) of deuterated 50mM sodium phosphate (pH 7) buffer with 2mM of ethylene diamine tetra-acetic acid (EDTA). The remaining 10% (v/v) of the sample was the internal standard (5mM DSS-D6 and 0.2% NaN3 in D2O; Chenomx Inc., Alberta, Canada). The final volume for the cell extract samples was 60μL (in 1.5 mm tube); the final volume for supernatant samples was 200μL (in 3 mm tube). To the J-resolved heteronuclear single quantum coherence spectroscopy (j-HSQC) ^12^ samples, [1,2-^13^C2] and [2-^13^C] acetate was added as standard in a 2:1 ratio, respectively.

### NMR spectroscopy

Nuclear magnetic resonance (NMR) spectra were collected at 14.1 T. ^13^C 1D conventional spectra were collected with a home-built superconducting (HTS) ^23^ probe. Both 1D proton and J-HSQC spectra (j-HSQC) were collected with a CP TXI CryoProbe and an Avance II Console (Bruker Biospin, Billerca, MA). Proton spectra were acquired using the first slice of a NOESY pulse sequence (tnnoesy) with a relaxation delay (d1) of 3 s, spectral width (sw) of 7211.54 Hz, mixing time of 100 ms, number of scans (nt) of 64, and acquisition time of 4 s. jHSQC spectra were collected with d1 of 1 s, 32 to 272 scans (depending on the sample mass), sw of 7211.54 Hz in f2 dimension and 150.93 Hz in f1 dimension, and an acquisition time of 0.20 s with GARP4 ^13^C decoupling. Conventional ^13^C spectra were acquired with 1.5 s d1, 240 ppm sw, and 1.4 s acquisition time, with proton decoupling using WALTZ-16. All experiments were acquired at room temperature (25°C).

### NMR Data analysis

All NMR spectra were processed using either MestReNova 11.0.0-17609 (Mestrelab Research, S.L., and Santiago de Compostela, Spain) or TOPSPIN 3.5 (Bruker, Billerica, MA, USA). Proton spectra were calibrated by setting the DSS singlet resonance to 0 ppm and were processed using a zero filling of 131072 points, 0.5 Hz exponential line broadening, and Whittaker-Smoother or spline baseline correction as required. Online-based Bayesil software^24^ (Chenomx NMR Suite 8.2 (Chenomx, Inc., Edmonton, Alberta, Canada)) and/or the Eretic method^25,26^ were employed to analyze concentrations of metabolites quantitatively. The Eretic method was utilized for some supernatant samples where the DSS peak broadened by the presence of macromolecules (proteins or lipids).

jHSQC spectra were processed using a quadratic sine function on both dimensions (Figure 5C and 5D). A shifted sine bell (SSB) factor was applied to produce spectra with less overlap at the foot of the peaks in the indirectly detected dimension. Sine bell shift (SSB) of 2.6 and 2 were applied in F2 and F1 dimension, respectively. Complex data was not acquired in the second dimension, as the J-resolved spectra are inherently symmetric. The spectra were symmetrized in mNOVA to further reduce artifacts in the 2D spectra. Calibration of chemical shifts was done by setting the acetate standard peak to 1.91 ppm. One dimensional projections of the carbon 4 (C4) resonance of glutamate was extracted from jHSQC spectrum. The peak multiples were fitted to a mixed Gaussian/Lorentzian shape to extract peak areas, and those values were input into the flux analysis^27^. ^13^C NMR spectra were processed with 0.5 Hz exponential line broadening and spline baseline correction. They were zero-filled to 131072 points.

### T1D incidence

All mice used in this study were housed in specific pathogen-free facilities as previously described^20^. These studies were approved by the Institutional Animal Care and Use Committee (IACUC) at The Jackson Laboratories and University of Florida. All mice were placed onto water bottles that contained either filter-sterilized water or filter-sterilized water containing 35mM or 70mM 2DG. Development of spontaneous T1D in control or 70mM treated NOD.AI4 α/β-Thy1^a/b^ (NOD.Ai4) mice was assessed from 3 weeks of age, and in 35mM treated NOD/ShiLtJ mice from 10 weeks of age. Diabetes development was monitored every other day starting from time of treatment. Incidence was determined using measurement of blood glucose from the tail vein using a One-Touch Ultra 2 Meter (Life Scan, Milpitas, CA). A blood glucose level >600mg/dl on two consecutive days was considered diagnostic of diabetes onset for NOD.Ai4 mice, and >200mg/dL for NOD/ShiLtJ mice.

### Flow cytometry assessment of immune cell phenotype

Flow cytometry analyses of the spleens of control and 2DG treated NOD/ShiLtJ mice, and peripheral blood and spleens of control and 2DG treated NOD.Ai4 mice, were carried out as previously described^20^. Blood samples were collected using EDTA coated tubes, and lymph nodes and spleens homogenized. Red blood cells were lysed using ACK lysis buffer (Corning, USA), and remaining immune cells stained with antibodies against CD45R, CD3, CD8a, CD44, CD19, CD73, Foxp3, Tcr-β, CD25, CD62L, and CD69. Tetramer, H-2D b tetramer loaded with mimetope (YAIENYLEL), or irrelevant lymphocytic choriomeningitis virus–derived peptide (FQPQNGQFI) loaded control H-2D b tetramer was used (both produced by the National Institutes of Health Tetramer Core Facility, Atlanta, GA) to ensure gated populations were insulin-reactive CD8^+^ T-cells. Flow cytometry was performed using a BD LSR Fortessa or BD FACSCalibur. Data were analyzed using FlowJo software (TreeStar, Ashland, OR). Antibody sources and clones detailed in **Supplemental Table 2**.

### Adoptive Transfer

10-week-old female NOD/ShiLtJ were treated for 10 weeks with 35mM 2DG in their drinking water or placed on control water bottles prior to adoptive transfer. Splenocytes from NOD/ShiLtJ mice were isolated as previously described^20^. After isolation, splenocytes were pooled, washed, and resuspended in ice-cold Dulbecco’s PBS at a concentration of 100 × 10^6^/mL. Cells (8.8 × 10^6^) were injected into NOD.CB17-Prkdcscid/J recipients *via* the tail vein. Mice were monitored for diabetes incidence and sacrificed upon diabetes onset.

6-week-old NOD/ShiLtJ female mice were treated with 70mM 2DG in their drinking water or placed on control water bottles for 2 weeks prior to adoptive transfer. Splenocytes from NOD.Ai4 mice were isolated as previously described^20^ and stained with CellTrace Violet (CTV) cell proliferation dye (ThermoFisher Scientific, USA), according to the manufacturer’s instructions. After labeling, splenocytes were washed and resuspended in ice-cold Dulbecco’s PBS at a concentration of 10 × 10^6^/mL. Cells (2 × 10^6^) were injected into sub lethally irradiated control and 2DG treated NOD/ShiLtJ female recipients via the tail vein. Three days post-transfer, mice were sacrificed, and pancreatic draining lymph nodes (PLNs), spleen, and peripheral blood collected. Spleens and lymph nodes were homogenized to single cells and stained for flow cytometry analyses. Cells were analyzed using Live/Dead near IR (ThermoFisher Scientific, USA) to exclude dead cells. NOD.Ai4 CD8^+^ T-cells were identified with anti-CD8 (BioLegend, San Diego, CA) and tetramer staining. Activation status of AI4 cells was determined by staining with antibodies against Ki-67. Data were analyzed using FlowJo software.

### Pancreatic Image Analysis

After euthanasia, the pancreata of control and 35mM 2DG treated NOD.Ai4 mice were fixed in 10% buffered formalin overnight, paraffin embedded, and stained with chromogen, anti-insulin (Agilent, USA), and anti-CD3 (Serotec, USA). Sections were mounted onto glass slides and scanned into Aperio eSlide Manager (Leica Biosystems, USA). In-built macros in the Aperio Imagescope (Leica Biosystems, USA) software were calibrated and used to analyze insulin and CD3 positive staining in whole pancreatic sections for both control and 35mM 2DG treated NOD.Ai4 mice.

### Statistical analysis

All statistical analyses were carried out using Prism 8 software (GraphPad, USA). Statistical tests were conducted by using non-parametric One-way ANOVA. Statistical significance was defined as *p<0.05*.

## Acknowledgments

These studies were supported by grants from the National Institutes of Health UC4 DK104197 (CEM), R01DK074656 (CEM), UG3DK122638 (CEM), P01 AI042288 (CEM), DK046266 (DVS), DK095735 (DVS), OD-020351-5022 (DVS), R01 DK105346 (MEM), U24 DK097209 (MEM), P41 GM122698 (MEM), F30 DK105788 (BNN), as well as by Juvenile Diabetes Research Foundation grant 2018-568, Department of Defense Grant T0048 (TEA), the University of Florida Experimental Pathology Innovative Grant (SES), and the Sebastian Family Endowment for Diabetes Research (CEM). The University of Florida Diabetes Institute and the University of Florida Center for Immunology and Transplantation supported these studies. A portion of this work was performed in the McKnight Brain Institute at the National High Magnetic Field Laboratory’s AMRIS Facility, which is supported by National Science Foundation Cooperative Agreement No. DMR-1644779 and the State of Florida. The authors acknowledge the assistance of Amanda Posgai and Sara Williams for review and formatting of the manuscript.

## Author Contributions

S.E.S., D.V.S., and C.E.M. conceived the study. S.E.S., T.E.A., M.M., D.V.S., and C.E.M. designed experiments. S.E.S., J.C., B.N.N., R.K., and H.D.C. performed the experiments. S.E.S., J.C., B.N.N., R.K., H.D.C., T.E.A., M.M., D.V.S., and C.E.M. analyzed the data. S.E.S. and C.E.M. and drafted the manuscript. All authors reviewed and contributed feedback on the final manuscript.

---

**Supplemental Figure 1.** *Early phenotypic changes observed in human naïve CD8^+^ T-cells in conditions of substrate limitation/excess after 4 days of activation*. **a-b)** Early activation and expansion in presence of metabolic inhibitions produced more memory-like cells compared to that of glucose alone. **c)** Early expression of activation marker, CD137, was not altered with glycolytic and mitochondrial inhibition. **d)** Decreased expression of cytolytic enzymes, Granzyme B, occurs in the presence of glycolytic and mitochondrial inhibition. **e)** Expression of CXCR3 is decreased when glycolytic and mitochondrial inhibition occurs. Mean ± SEM graphed. These data are compiled from 4 healthy donors. ᵻ denotes *p<0.05*, ᵻᵻ denotes *p<0.005 and* ᵻᵻᵻ denotes *p*<0.0001.

**Supplemental Figure 2.** *No alterations in naïve, memory, or effector CD8^+^ T-cell populations in 35mM 2DG administered NOD.AI4αβ mice*. Weekly cheek bleeds were carried out and naïve **a)** memory **b)** and effector **c)** CD8^+^ T-cells monitored in control and 35mM treated mice until diabetes onset was reached. Upon reaching diabetes onset, mice were sacrificed, and spleens and pancreatic lymph nodes harvested with naïve **d, g)** memory **e, and h)** and effector **f, i)** CD8^+^ T-cells populations analysed. Mean ± SD graphed.

**Supplemental Figure 3.** *70mM 2DG treatment does not alter naïve, memory, or effector CD8^+^ T-cell populations in NOD.AI4αβ mice*. Weekly cheek bleeds were carried out and naïve **a)** memory **b)** and effector **c)** CD8^+^ T-cells monitored in control and 35mM treated mice until diabetes onset was reached. Upon reaching diabetes onset, mice were sacrificed, and spleens and pancreatic lymph nodes harvested with naïve **d, g)** memory **e, and h)** and effector **f, i)** CD8^+^ T-cells populations analysed. Mean ± SD graphed.

**Supplemental Figure 4.** 2DG does not arrest CD8+ T-cell movement or interactions. Primary human islets were stained with TMRM (red) and embedded along with IGRP-GFP CD8+ T-cells (green) in Matrigel and imaged for 16 hours in the presence of either 5.5mM glucose **a)** or 2.7mM D-glucose plus 2.7mM 2DG **b)**. T-cell movement was tracked and velocity **c)** calculated, as well as displacement for control **d)** and 2DG **e)**, with no significant differences in the T-cells’ movement or velocity.

