## Supplemental Figures 1-4 for "Human CD8^+^ T-cells Require Glycolysis to Elicit Effector Function"

**Supplemental Table 1**. *Reporting of human islet preparation used*.

| **Islet Preparation** | **1** |
|---|---|
| **Donor demographics** | |
| Unique identifier | SAMN08912443 |
| Age (yrs) | 38 |
| Sex (M/F) | Male |
| BMI | 23.90 |
| Ethnicity | Hispanic/Latino |
| $HbA_{1c}$ | n/a |
| COD | Cerebrovascular/ Stroke |
| **Pancreas** | |
| Warm ischaemia time (h) | No |
| Cold ischaemia time (h) | 7 hours 57 minutes |
| **Islet handling and use** | |
| Origin/source | IIDP |
| Isolation centre | Southern California Islet Resources Center |
| Estimated purity (%) | 80% |
| Estimated viability (%) | 94% |
| Total culture time (hrs) | 19 hours |
| Functional measurement | |
| Islet handling | Islets handpicked; media changed into cDMEM |
| Handpicked to purity? | Yes |
| Experimental islet use (including in which experiment each islet preparation was used) | 3D culture experiments |

**Supplemental Table 2:** *Antibody sources and clones used in the investigation*. All antibodies were used as per manufacturer recommendations.

| ***Marker*** | ***Species Reactivity*** | ***Color*** | ***Clone*** | ***Company*** |
|---|---|---|---|---|
| *CD3* | Mouse | BV421 | 17A2 | BioLegend, USA |
| *CD4* | Mouse | APC | RM4-5 | BioLegend, USA |
| *CD4* | Mouse | PE | GK1.5 | BioLegend, USA |
| *CD8a* | Human | APC/Cy7 | SK1 | BioLegend, USA |
| *CD8a* | Human | PE | SK1 | BioLegend, USA |
| *CD8a* | Mouse | BV605 | 53-6.7 | BioLegend, USA |
| *CD19* | Mouse | FITC | 1D3 | BD Pharmingen, USA |
| *CD25* | Mouse | PE/Cy7 | HK1.4 | BioLegend, USA |
| *CD44* | Mouse | AF488 | IM7 | BioLegend, USA |
| *CD45RA* | Human | PE/Cy7 | HI100 | BioLegend, USA |
| *CD45R* | Mouse | BV510 | Ra3-6B2 | BioLegend, USA |
| *CD45RO* | Human | PerCP/Cy5.5 | UCHL1 | BioLegend, USA |
| *CD62L* | Human | APC | DREG-56 | BioLegend, USA |
| *CD62L* | Mouse | APC | MEL14 | BioLegend, USA |
| *CD69* | Human | PerCP | FN50 | BioLegend, USA |
| *CD69* | Mouse | PerCP/Cy5.5 | H1.2F3 | BD Pharmingen, USA |
| *CD73* | Mouse | PE | TY/11.8 | BioLegend, USA |
| *CD137* | Human | PE | 4B4-1 | BioLegend, USA |
| *CXCR3* | Human | BV510 | G02H7 | BioLegend, USA |
| *Foxp3* | Mouse | AF488 | MF23 | BD Pharmingen, USA |
| *Ki-67* | Human | AF700 | Ki-67 | BioLegend, USA |
| *Granzyme B* | Human | AF647 | GB11 | BioLegend, USA |
| *Tcr-β* | Mouse | FITC | H57-597 | BD Pharmingen, USA |

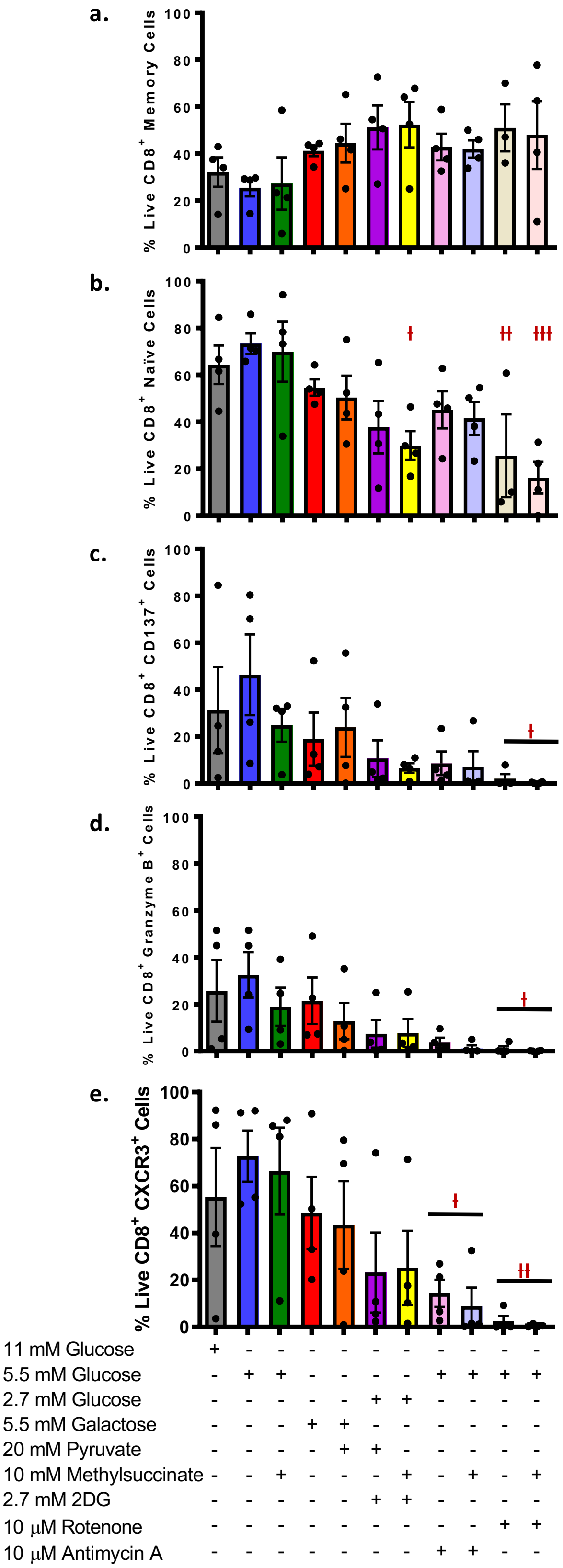


**Supplemental Figure 1.** *Early phenotypic changes observed in human naïve $CD8^+$ T cells in conditions of substrate limitation/excess after 4 days of activation*. **a-b)** Early activation and expansion in presence of metabolic inhibitions produced more memory-like cells compared to that of glucose alone. **c)** Early expression of activation marker, CD137, was not altered with glycolytic and mitochondrial inhibition. **d)** Decreased expression of cytolytic enzymes, Granzyme B, occurs in the presence of glycolytic and mitochondrial inhibition. **e)** Expression of CXCR3 is decreased when glycolytic and mitochondrial inhibition occurs. Mean ± SEM graphed. These data are compiled from 4 healthy donors. ł denotes *p<0.05*, łł denotes *p<0.005 and* łłł denotes *p<0.0001*

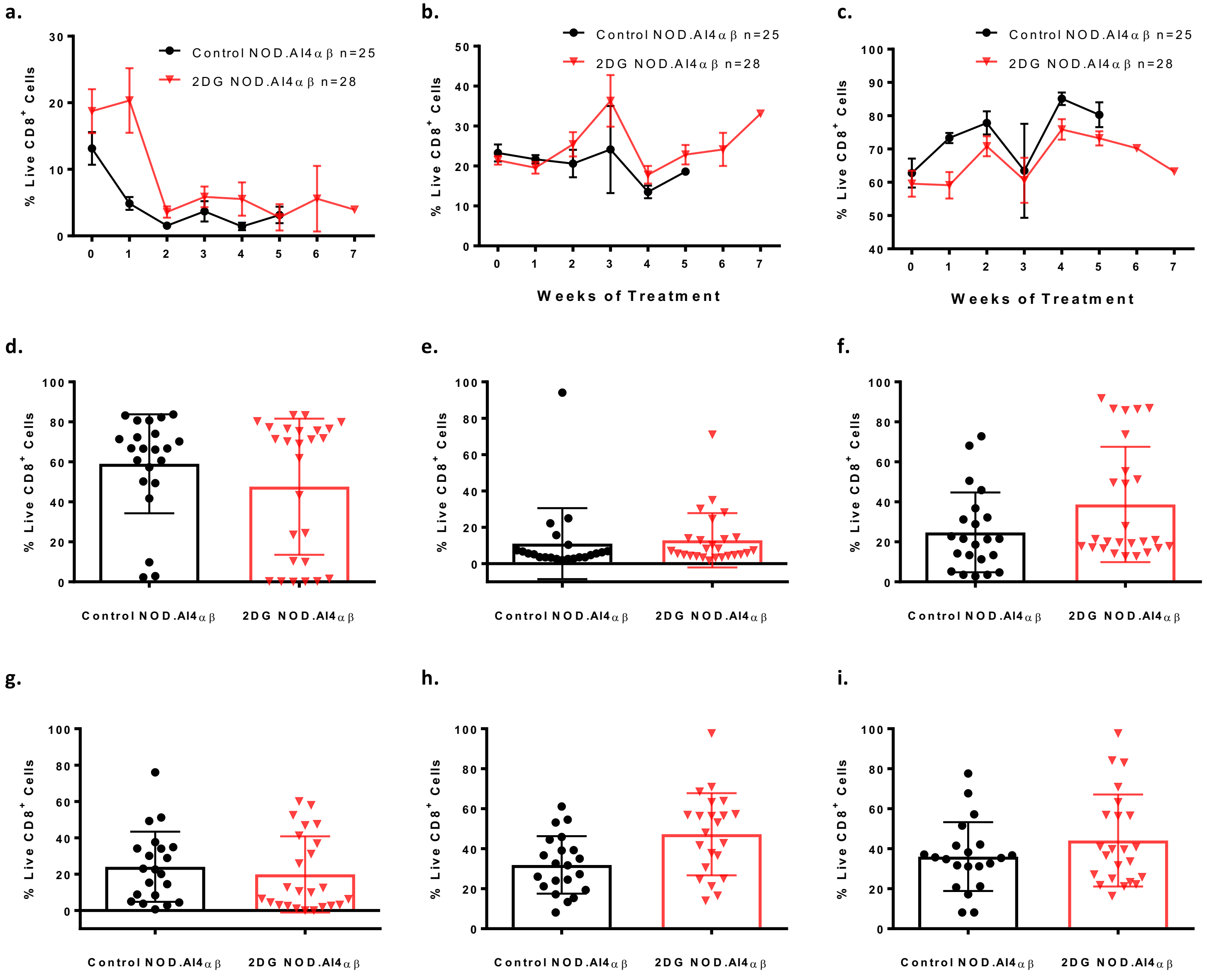


**Supplemental Figure 2.** *No alterations in naïve, memory or effector CD8⁺ T cell populations in 35mM 2DG administered NOD.AI4αβ mice*. Weekly cheek bleeds were carried out and naïve **a)** memory **b)** and effector **c)** CD8⁺ T cells monitored in control and 35mM treated mice until diabetes on-set was reached. Upon reaching diabetes on-set mice were sacrificed and spleens and pancreatic lymph nodes harvested with naïve **d,g)** memory **e,h)** and effector **f,i)** CD8⁺ T cells populations analysed. Mean ± SD graphed

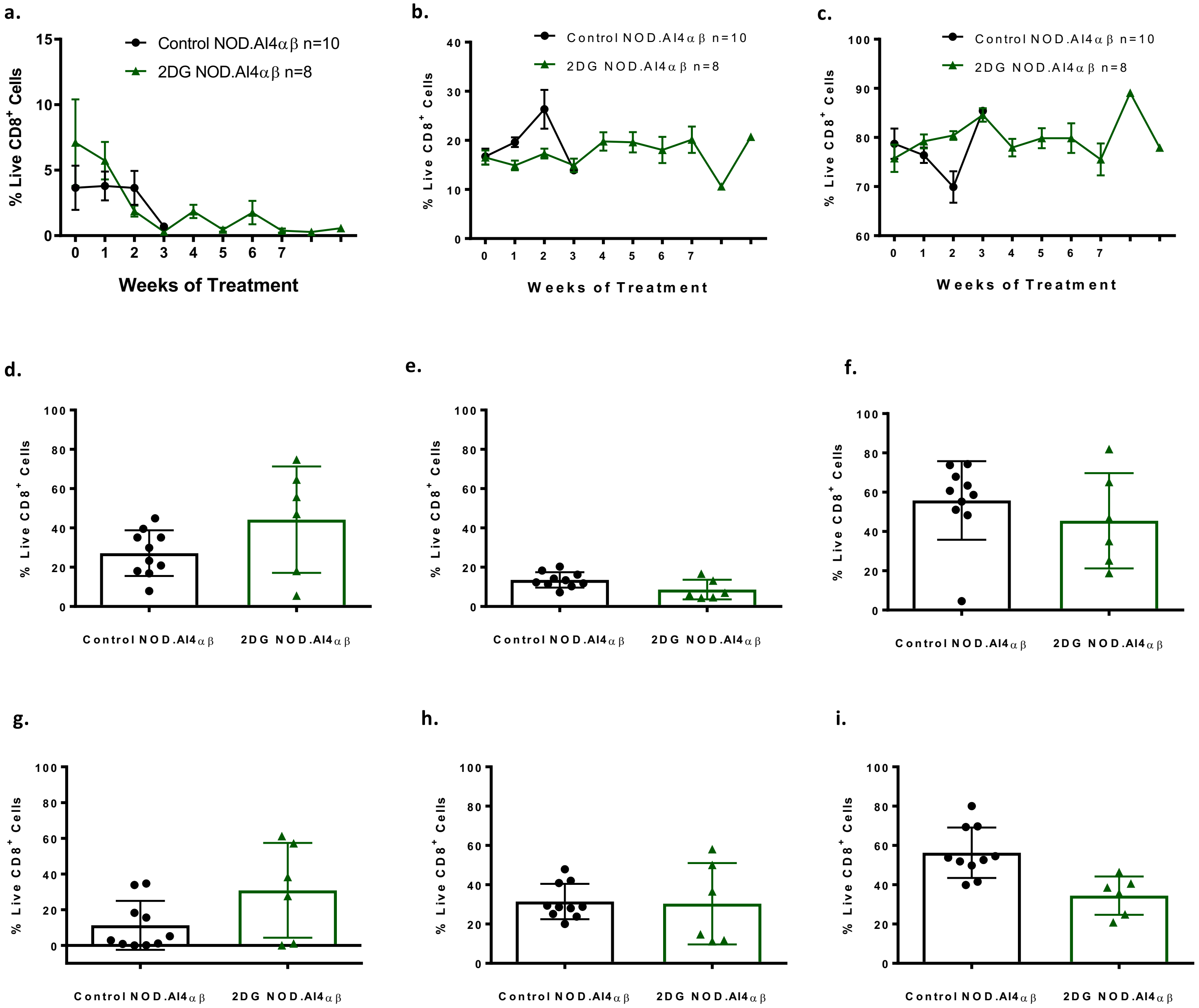


**Supplemental Figure 3.** *70mM 2DG treatment does not alter naïve, memory or effector CD8⁺ T cell populations in NOD.AI4αβ mice*. Weekly cheek bleeds were carried out and naïve **a)** memory **b)** and effector **c)** CD8⁺ T cells monitored in control and 35mM treated mice until diabetes on-set was reached. Upon reaching diabetes on-set mice were sacrificed and spleens and pancreatic lymph nodes harvested with naïve **d,g)** memory **e,h)** and effector **f,i)** CD8⁺ T cells populations analysed. Mean ± SD graphed.

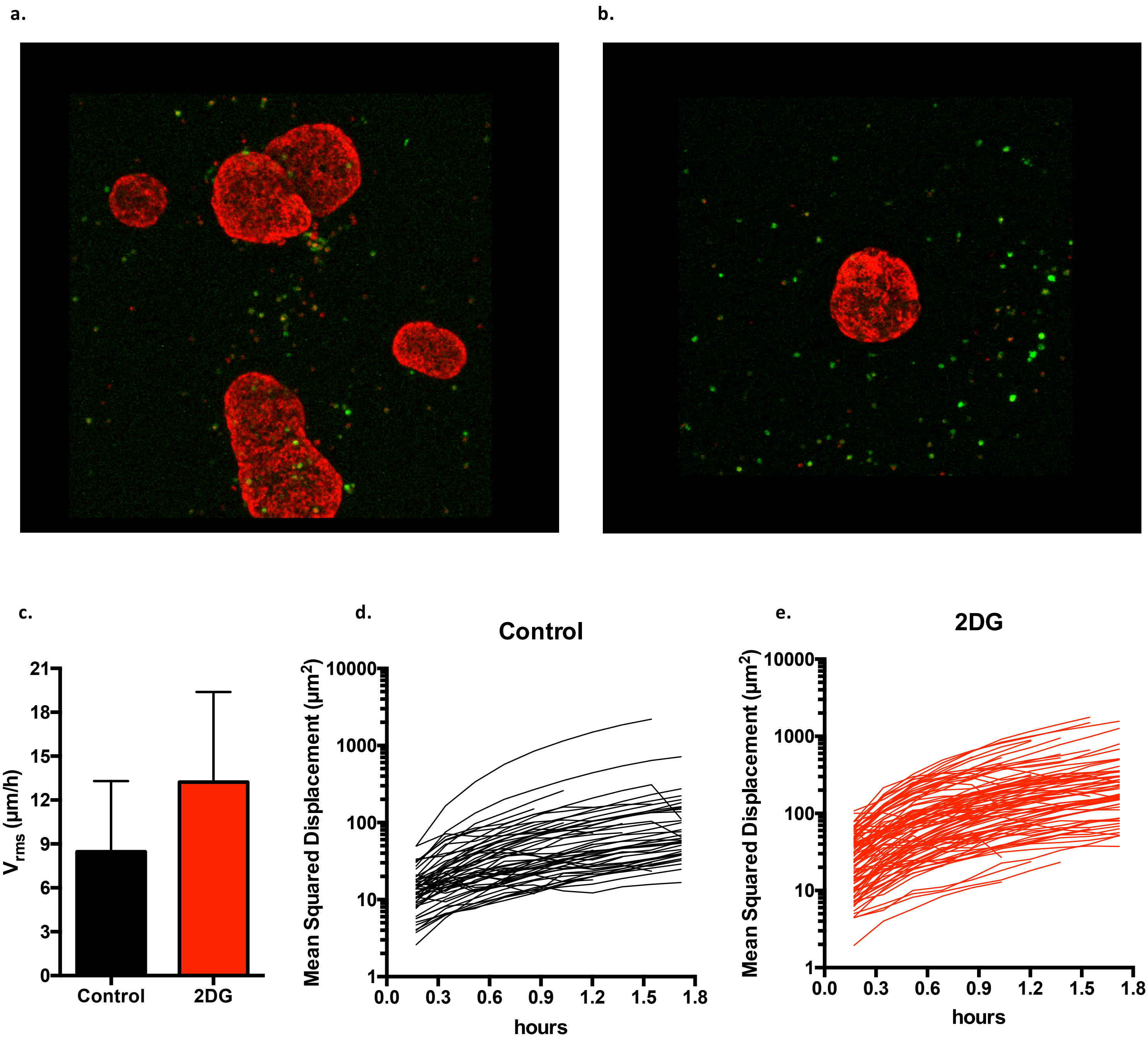


**Supplemental Figure 4.** *2DG does not arrest CD8⁺ T cell movement or interactions*. Primary human islets were stained with TMRM (red) and embedded along with IGRP-GFP CD8⁺ T cells (green) in Matrigel and imaged for 16 hours in the presence of either 5.5mM glucose **a)** or 2.7mM D-glucose plus 2.7mM 2DG **b)**. T cell movement was tracked and velocity **c)** calculated as well as displacement for control **d)** and 2DG **e)**, with no significant differences in the T cells movement or velocity.
